# Macroevolutionary integration of phenotypes within and across ant worker castes

**DOI:** 10.1101/604470

**Authors:** Nicholas R. Friedman, Beatrice Lecroq Bennet, Georg Fischer, Eli M. Sarnat, Jen-Pan Huang, L. Lacey Knowles, Evan P. Economo

**Affiliations:** Okinawa Institute of Science and Technology Graduate University, 1919-1 Tancha, Onna-son, Okinawa, Japan 904-0495; Museum of Zoology, Department of Ecology & Evolutionary Biology, University of Michigan; Biodiversity Research Center, Academia Sinica, Taipei 11529, Taiwan

**Keywords:** Morphological integration, modularity, caste, dimorphism, *Pheidole*, ants, geometric morphometrics

## Abstract

Phenotypic traits are often integrated into evolutionary modules: sets of organismal parts that evolve together. In social insect colonies the concepts of integration and modularity apply to sets of traits both within and among functionally and phenotypically differentiated castes. On macroevolutionary timescales, patterns of integration and modularity within and across castes can be clues to the selective and ecological factors shaping their evolution and diversification. We develop a set of hypotheses describing contrasting patterns of worker integration and apply this framework in a broad (246 species) comparative analysis of major and minor worker evolution in the hyperdiverse ant genus *Pheidole*. Using geometric morphometrics in a phylogenetic framework, we inferred fast and tightly integrated evolution of mesosoma shape between major and minor workers, but slower and more independent evolution of head shape between the two worker castes. Thus, *Pheidole* workers are evolving as a mixture of intra- and inter-caste integration and rate heterogeneity. The decoupling of homologous traits across worker castes may represent an important process facilitating the rise of social complexity.

## Introduction

The increase of morphological complexity following divergence in cellular function is a repeating theme in the evolution of multicellular organisms (Wagner and Altenberg 1996). Given cues regarding their developmental fate, cells and tissues express their identical genomes in different ways to produce different traits and thus allow functional specialization. Morphological integration can be considered the extent to which these traits vary in concert, either as a continuation of their shared genetic or developmental origin, or as a unification of parts contributing to a shared function and shaped by selection (Olson and Miller 1958; Klingenberg 2008). Sets of integrated traits covary as modules, between which covariation is weaker than within (as in the primate cranium; Cheverud 1982).

Much as a single genome can underlie different cooperating tissues and traits within the same organism, different traits are also produced among individuals using the same genome. Distinct phenotypes are commonly observed in different sexes (Owens and Hartley 1998), or in individuals adopting alternative reproductive tactics (Emlen et al. 2007) as a result of differential selection. Eusocial insects reflect a major evolutionary transition whereby a unit of selection is comprised of different individuals working together as part of an integrated colony-level phenotype (Wheeler 1911; Hölldobler and Wilson 1990; Szathmáry and Smith 1995) and understanding the evolution and function of these “superorganisms” is a major and enduring interest of evolutionary biology (Oster and Wilson 1978; Seeley 1995; Holldobler and Wilson 2009). The castes of social insects can exhibit radically different traits from the same genome: a female egg laid by the queen has the potential to develop into either another queen or a worker caste individual. This phenotypic polymorphism allows functional specialization among individuals in a colony and the rise of social complexity, the feature of eusociality that best defines its potential for division of labor (Oster and Wilson 1978; Hölldobler and Wilson 1990). While worker castes are an ancestral trait shared by nearly all extant ants, several lineages have since evolved further division of labor among workers to form worker castes – known also as subcastes (Wilson 1953; Hölldobler and Wilson 1990; Oster and Wilson 1978; Wills et al. 2017). In the colonies of some species (e.g., *Solenopsis invicta)*, worker castes exhibit polymorphism mostly along a single allometric function – shape varies with size along a regular continuum (Wilson 1953). However, for species in other genera (e.g., *Pheidole, Colobopsis, Carebara, Cephalotes, Eciton, Acanthomyrmex, Pseudolasius)*, variation reaches “complete dimorphism” into distinct major worker and minor worker phenotypes (Wilson 1953). While there is some contention over nomenclature within the myrmecological community (Urbani 2015), we refer to minor workers and major workers (aka soldiers) as “worker castes” throughout following Wills et al. (2017).

The evolution of complete dimorphism offers the potential for new dimensions of variation in ants (Wilson 1953, Wills et al. 2017). If phenotypes are disintegrated among worker castes, this can allow for greater functional specialization and different combinations of traits available to the colony-level phenotype (Wilson 1953; Powell 2008; Powell 2009; Wills et al. 2017). However, the evolution of specialized morphology in major workers may be biased by developmental pathways that are shared with minors (Wheeler and Nijhout 1983; Wheeler and Nijhout 1984, Wheeler 1991; Rajakumar et al. 2012), thus there could be limits to divergence among homologous body parts across the different worker castes, or a shared pathway could lead selection on one worker caste to result in a neutral change in the other.

The ecological and behavioral roles of polymorphic worker ants have long been a fascination of social insect research (Wheeler 1911; Goetsch 1937; Wilson 1953, Oster and Wilson 1978; Powell and Franks 2006; Powell 2008; Powell 2009; Powell 2016; Wills et al. 2017). Likewise, the genomic and biochemical mechanisms underpinning caste differentiation is a central avenue for understanding the evolution of social complexity (Wheeler 1991; Hughes et al. 2003, Anderson et al. 2008; Molet et al. 2012; Rajakumar et al. 2012; Lillico-Ouachour and Abouheif 2017, Gospocic et al. 2017, Chandra et al. 2018). However, the macroevolutionary implications of these processes—the patterns of integration and modularity that emerge across the diversification of hundreds or thousands of lineages—are comparatively less well-studied in the ants, although the topic is receiving increasing interest (Pie and Traniello 2007; Pie and Tschá 2013; Holley et al. 2016; Powell 2016). These patterns, revealing the degree to which different traits evolve independently within and among worker castes, may be an important clue to both the selective forces driving evolution like ecological subspecialization among worker castes (Powell and Franks 2006), and the potential constraints on evolution like restrictive developmental limitations (as in Fritz et al. 2014). Furthermore, patterns of evolutionary rate heterogeneity or homogeneity within and across worker castes may reflect particular aspects of the phenotype that are under strong selection because they underlie axes of ecological divergence among species (Schluter 2000; Price et al. 2016).

Ants have colonized and evolved adaptations to many environments, and are among the most abundant terrestrial organisms on the planet. Ants have also radiated to produce a diverse array of morphologies in nearly every region they have colonized (Hölldobler and Wilson 1990). In this study, we focus on overall body size, the relative body sizes of different parts, and the shapes of the head and mesosoma. Body size has been shown to be a major axis of morphological variation among ants (Pie and Traniello 2007, Price et al. 2016; Powell 2016). Previous studies of functional morphology in ants have also focused on the head (e.g., Holley et al. 2016), which contains the main apparatus for feeding (mouthparts, mandibles), manipulating objects (mandibles), and sensation (eyes, antennae). If head shape is the primary focus of ecological adaptation, this trait should evolve more rapidly than others during radiation. We also examine the mesosoma, the main power center of the ant including muscles for bearing loads and moving the legs. While the functional significance of external mesosoma shape is not well understood, the shapes and relative sizes of different regions likely reflects investment in different muscle groups that have functional implications. The sizes and positions of the sclerites (plates of the cuticle which are partially captured here by our landmarks) are associated with homologous attachment points underneath. For example, Keller et al. (2014) showed that the pronotal region associated with the T1 sclerite houses the muscles that lift the head. Other regions of the mesosoma contain stabilizing muscles, muscles to support the legs, and muscles to flex the petiole (Lubbock 1881), all of which have obvious functional implications.

Relatively few studies have compared the tempo of evolution across different ant traits, (but see Pie and Tschá 2013; Blanchard and Moreau 2017; Holley et al. 2016). If the shapes of other traits such as the mesosoma (thorax) evolve more rapidly, this may be an indication that they serve a greater functional role in ecological divergence than previously understood. Likewise, if majors exhibit greater rates of change, that may signal that their functional role has changed often following the evolution of complete dimorphism, or that they are important for achieving and maintaining ecological divergence among species.

To compare morphological integration and evolutionary rate of different worker castes and traits, we focused on the ant genus *Pheidole*. The ants of this genus have, in the course of their approximately 37 million year history, spread throughout 6 continents to produce more than 1000 described (and many more undescribed) species (Moreau 2008; Economo et al. 2015a). Perhaps the most notable characteristic of species in this hyperdiverse genus is the clear dimorphism of their workers: a major worker caste with enlarged heads is easily visible in all species (indeed a third super-major form is also observed in some species) (Wilson 2003). Behavioral studies have described different ecological roles for *Pheidole* worker castes, with major workers performing more defense, food processing, and storage tasks than minor workers (Wilson 1984; Tsuji 1990; Mertl and Traniello 2009; Huang 2010). The relatively consistent body plan and caste structure of this genus make it an ideal clade for comparative studies of morphology (Pie and Traniello 2007; Holley et al. 2016). The developmental basis of worker caste differentiation in *Pheidole* has been well studied over the years (Wheeler and Nijhout 1983; Wheeler and Nijhout 1984; Rajakumar et al. 2012; Lillico-Ouachour and Abouheif 2016; Rajakumar et al. 2018), and recent work on the taxonomy, biogeography, and ecomorphology of this group (Wilson 2003; Mertl and Traniello 2009, Muscedere and Traniello 2012; Sarnat and Moreau 2011, Economo and Sarnat 2012, Economo et al. 2015b; Holley et al. 2016; Sarnat et al. 2017) make it an attractive model clade for evolutionary research on social insects.

Several previous studies on the macroevolution of *Pheidole* morphology are particularly relevant for the current investigation. First, in an analysis before a *Pheidole* phylogeny was available, Pie and Traniello (2006) analyzed morphology with linear measurements and found that size differences explained most of the variation in *Pheidole* morphology across species, but majors and minors showed divergent patterns of character correlation. Later, with the benefit of a *Pheidole* phylogeny (Moreau 2008), Pie and Tschá (2013) showed that size varied more quickly than shape variables based on linear morphometrics, but did not explicitly test for modularity and integration. Holley et al. (2016) found that known ecological specialization of majors (seed milling behavior in granivorous species) was related to divergence in head size between major and minor worker castes (although enigmatically, due to a change in the minors), evidence that independent evolution of the two worker castes in relation to ecology can occur. Finally, Sarnat et al. (2017) tested hypotheses for the evolution of exaggerated thoracic spines, an unusual and geographically restricted phenotype in *Pheidole*.

Despite the insights of these pioneering studies, a comprehensive picture of the roles of integration, modularity, and rate heterogeneity in morphological evolution within and among *Pheidole* castes has not emerged. Using landmark-based geometric morphometrics, and taking advantage of recent progress on reconstructing the *Pheidole* phylogeny (Economo et al. 2015a; Economo et al. 2019) which allows for a more taxonomically and geographically extensive analysis, we perform the most morphologically and phylogenetically comprehensive analysis to-date to attempt to infer a general picture of integration and modularity in size and shape in the *Pheidole* worker castes.

To frame our study, we propose a set of hypotheses predicting different patterns of morphological integration within and among castes in social insect colonies (see Figure 1). We discuss this in terms of the head and mesosoma (thorax) of *Pheidole* worker castes, but it could equally be applied to any morphological traits shared among castes, or indeed traits shared among other differentiated phenotypes like sexes or reproductive strategies (Simpson et al. 2011). First, different parts of the body *within* a worker caste may be more or less integrated. This integration could reflect developmental biases or biomechanical constraints, for example a specific change in head morphology may necessitate a specific change of the thoracic segments that support or move the head. Second, *across* worker castes the same homologous body parts could be more or less integrated. As different worker castes share not only genomes but developmental pathways, it is plausible that selection on a trait in one worker caste could lead to a change in another worker caste. For example, selection on elongation of the head of a minor worker may lead to similar elongation in the major worker, even if there is no fitness benefit to the change in the major worker. Or, each worker caste could vary independently facilitating different functional roles in the colony.

**Figure 1:**
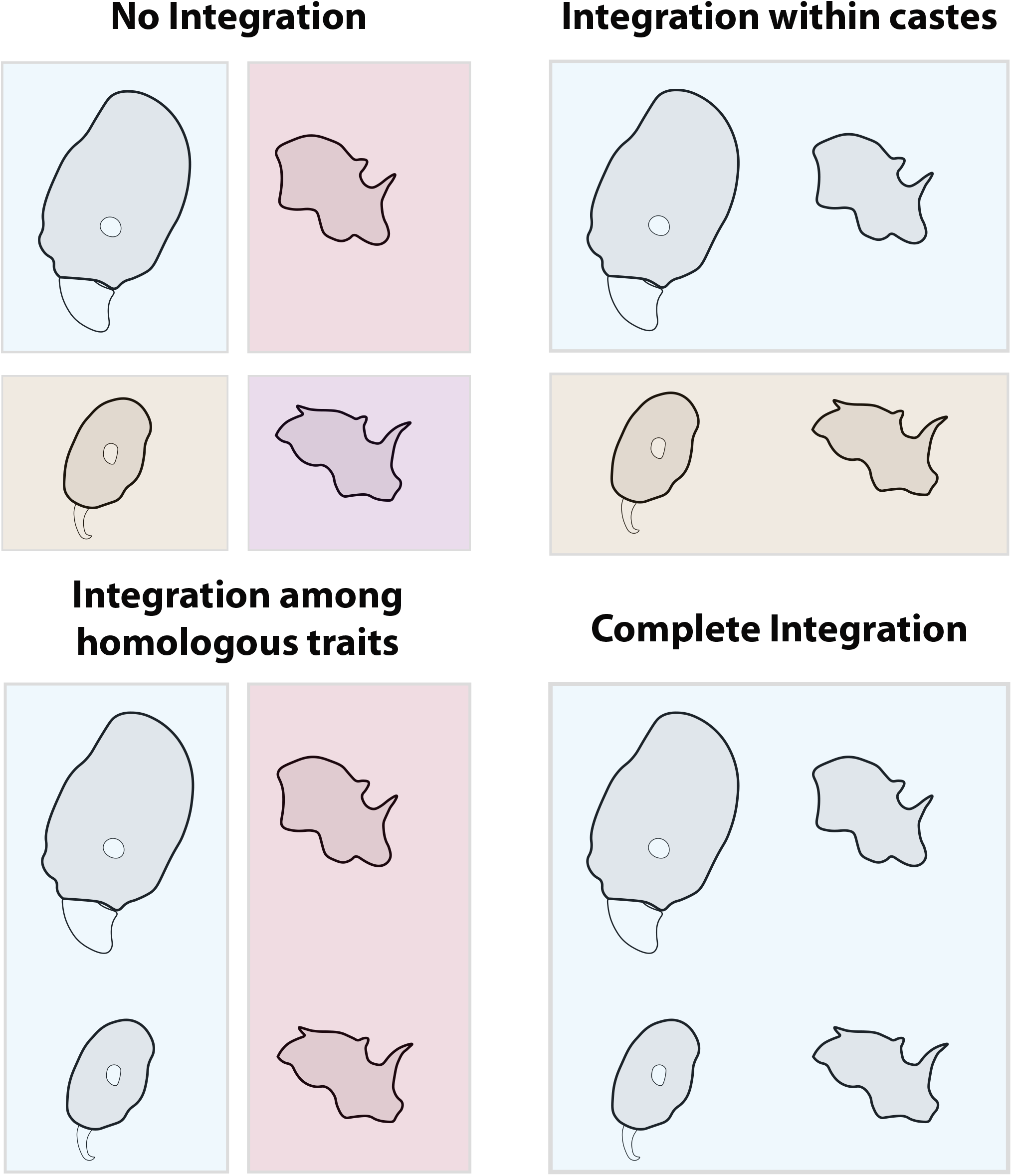
Hypothesized scenarios for the evolution of differentiated phenotypes. Worker castes or body parts united in the same box represent a pair of integrated traits. The scenarios we propose can be arranged in order of their extent of integration among homologous traits in different castes and among different traits within a caste.

We test these hypotheses by assessing the presence and pattern of integration of the head and mesosoma within and among worker castes. First, we assess heterogeneity in rates of evolution across body parts and worker castes; whether evolutionary change tends to follow a pattern in which different parts or worker castes are hot or cold spots of change, or whether there is rate homogeneity within and among worker castes. Second, we look for patterns of modularity in shape and size to test how well an evolutionary change in shape or size of one trait predicts the shape and size of another trait within the same worker caste or in a different one. If there is rate variation, we ask again whether those differences reflect characteristics shared among homologous traits or among worker castes.

## Methods

### Photographic Measurements

All comparative studies reflect a compromise between depth of individual sampling within species versus breadth across species. In this study we aimed to expand the latter to include as many *Pheidole* species as possible. We acknowledge a drawback of this strategy, which is that we cannot capture the size or shape range of individuals within each species. We measured a total of 1164 specimens from 314 species, measuring an average of 2.18 major worker and 2.20 minor worker specimens per species (Appendix S1); to maintain consistency between samples, all measurements were performed by coauthor BL. Myrmecologists use high resolution montage photographs to document ant diversity, following a standardized set of specimen positions that display head and body features from a consistent angle as described by the online resource and repository, AntWeb.org. We made a broad effort to photograph specimens from species used in recent phylogenetic projects (Economo et al. 2015a), supplemented with photographs taken by others and deposited on Antweb.org. We endeavored to collect data on both major and minor workers whenever possible, however photographic data for both worker castes were only available for 214 species or 68% of our total taxonomic sample. To account for potential focal length issues when using 2D photographs taken with different optical systems, we landmarked the same specimen 100 times under six different magnifications. A focal length warping effect was observable but was non-significant, and was within the range of intraspecific variation.

For each specimen, we placed landmarks using the three standard photographic angles: head view, dorsal view, and profile view. We collected landmarks from features that were consistently in the plane of the camera angle. Specifically, we placed 11 landmarks on the dorsal view of the head (Table S1) and 6 landmarks on the profile view of the body (Table S2; all located on the mesosoma; hereafter head, mesosoma; see Figure 2). To capture information on the posterior head shape, we also included a set of 6 sliding semi-landmarks (7 in major workers) from landmark 3 to 11 (Figure 2). The landmarks on the left side of the head were reflected bilaterally to produce the curve on the right side of the head between landmarks 11 and 1. Fixed landmarks on opposite sides of the head were reflected and averaged to force object symmetry.

**Figure 2:**
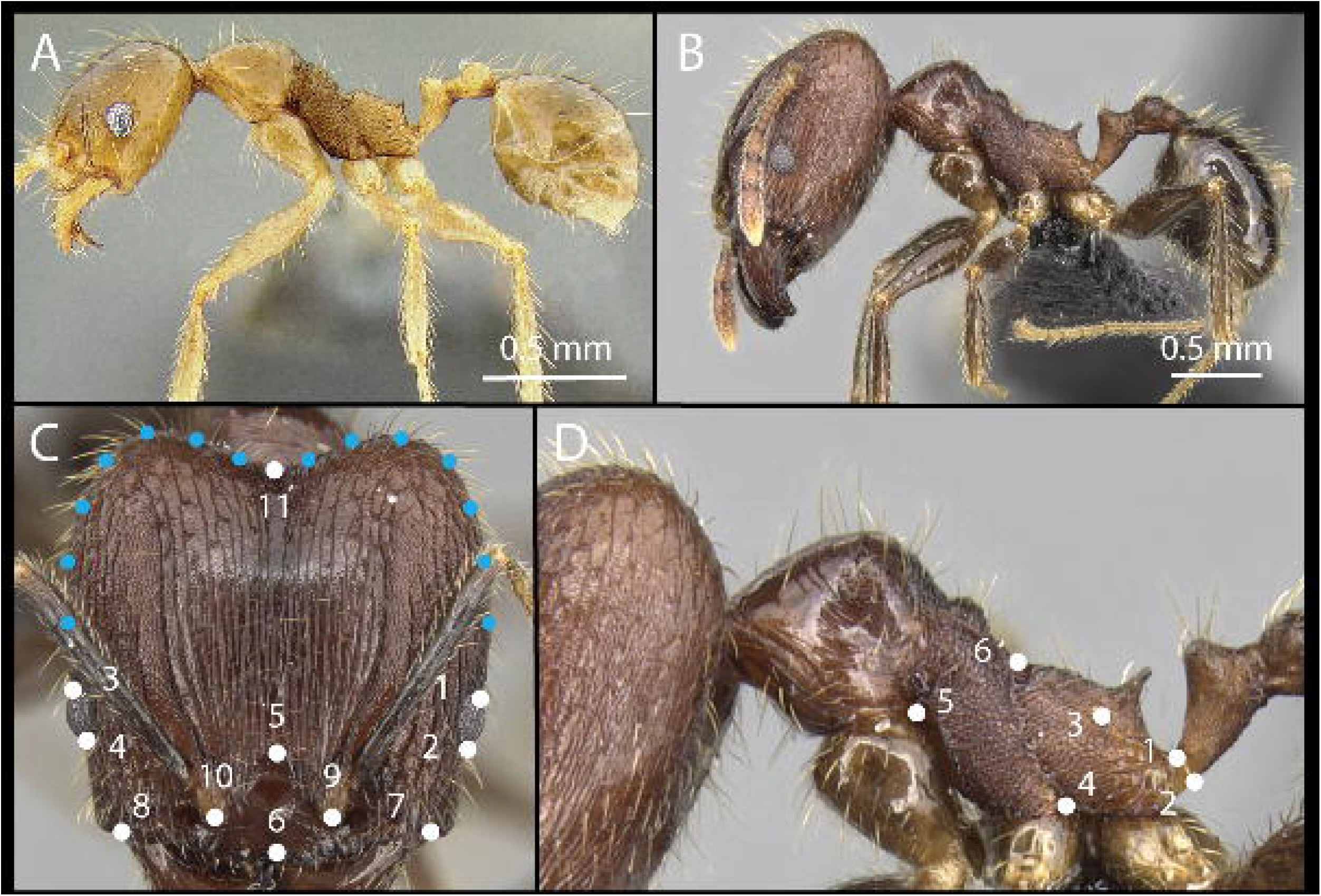
Example photographs of *fervens* minor worker (A) and major worker (B) assembled by photo-montage according to AntWeb specifications. Landmarks, in white, were placed on homologous features on the head (C) and mesosoma (D). Semi-landmarks, in blue, were spaced equally on the left side of the head between landmarks 3 and 11, and between landmarks 11 and 1.

While these landmarks omit several features that vary among *Pheidole* taxa, and those typically used in myrmecology research and taxonomy (Pie and Traniello 2007), this was unavoidable due to the constraints of choosing homologous landmarks in positions that are not occluded by nearby features (e.g., the anterior pronotum is often occluded by the posterior head lobes).

### Geometric Morphometrics

We performed a generalized Procrustes alignment on each set of landmarks using the R package *geomorph*, employing separate analyses for major and minor workers (Adams and Otárola-Castillo 2013; version 3.0.7). Specimens showing greater than expected distance from the Procrustes mean (i.e., above the upper quartile) were inspected for improper scale entry or landmark order/placement. Photos for which improper specimen positioning was observed were removed from the data set (< 1% of specimens studied). Within each species, we calculated the average Procrustes shape before proceeding with further analyses; we also averaged linear measurements in this manner. To visualize variation in highly dimensional shape characters, we estimated principal component axes and plotted species averages in tangent space (Figure 3c and e). As a proxy for body size, we used the logarithm of the centroid size of mesosoma landmarks as in (Economo et al. 2015a), which behaves similarly to the Weber’s Length measurement typically used by myrmecologists (Weber 1938). Only multivariate Procrustes alignment data, and not principle component data, were used in the comparative methods below (Uyeda et al. 2015).

**Figure 3:**
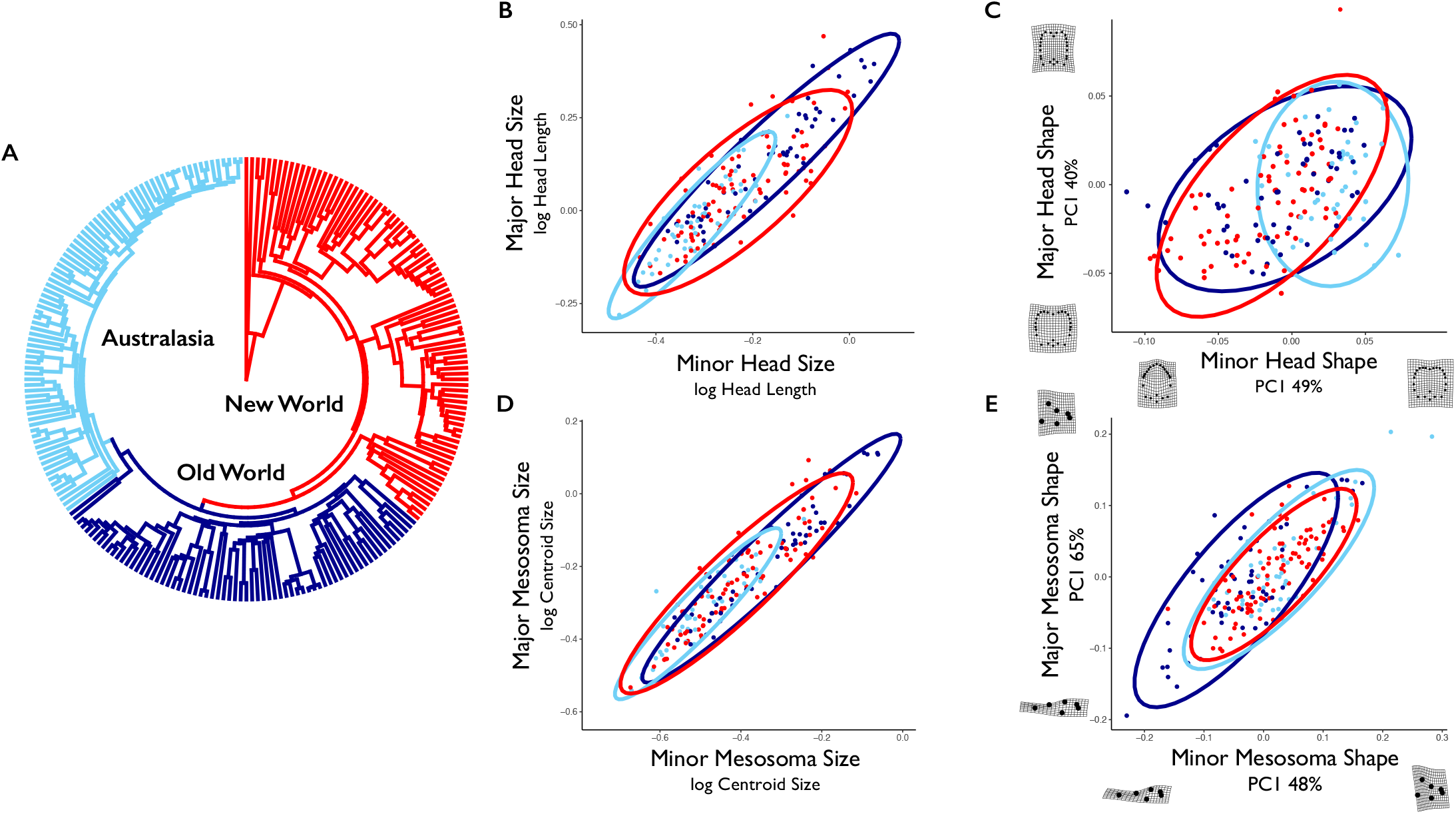
A phylogeny of the ant genus *Pheidole*, with clades colored by their geographic region, is shown in (A). Note that each clade represents a single colonization event (see Economo et al. 2015a). Comparisons of values for like traits in different castes are shown for head size (B), head shape (C), mesosoma size (D), and mesosoma shape (E). Ellipses reflect 95% confidence intervals, and are colored according to clade as in (A). For the shape data displayed in (C) and (E), the first principle component is shown for display purposes (and is not used in subsequent comparative methods), along with the percentage of variance it explains and deformation grids describing extreme values along the axis (produced using *geomorph;* Adams et al. 2018).

### Phylogenetic Data

We used a time-resolved phylogeny reconstructed by Economo et al. (2018) that includes 449 ingroup *Pheidole* species, based on a molecular dataset of nine loci. This phylogenetic tree builds upon previous analyses of *Pheidole* (Moreau 2008; Economo et al. 2015a), with the addition of 164 taxa and an expanded set of loci sequenced across species. For analyses in this paper, we used the maximum clade credibility tree from a Bayesian posterior set, which was pruned to contain only the taxa present in our morphological data (Figure 3; Figure S1).

### Comparative Methods

To examine the degree of correlated evolution between body regions (i.e., morphological integration), we used the *R* package *geomorph* (Adams and Otárola-Castillo 2013). We ran a series of pairwise integration tests between body regions and worker castes (Adams and Felice 2014). In each test, we estimated partial least squares (PLS) correlations between two sets of landmarks while correcting for phylogeny. The coefficient of correlation (r-PLS) for this regression describes the degree of integration. To calculate a p-value and significance test, we generated 1000 permutations of species’ phylogenetically-transformed values for each comparison. To compare integration of body size among worker castes and between the head and mesosoma, we used the coefficient of correlation for the linear regression of phylogenetically independent contrasts, hereafter r-PIC (Felsenstein 1985). To compare the relationship between trait shape and body size, we used a phylogenetic regression implemented for Procrustes shape variables (Adams and Collyer 2018). These and other methods described below were also run for the New World, Old World, and Australasian clades individually (Figure 3A; see online supplement). It is important to note that integration may exceed the values estimated here using PLS, as integration may span multiple PLS axis dimensions beyond the first axis, which is what we compared.

We used *geomorph* to estimate evolutionary rates for landmarked specimens (Denton and Adams 2015). As a significance test for differences in rates between traits, we performed 1000 simulations of trait evolution under a joint Brownian motion model, and compared the ratio of independently estimated rates to this simulated null. Given that differences in the number of landmarks can bias the amount of variation and thus rate described by each trait (Denton and Adams 2015), we report rate ratios for each pair of traits (e.g., major head vs. major mesosoma) as a proportion of the simulated null ratio.

We tested for evidence of evolutionary modularity within each body region (i.e., in addition to the head and mesosoma) again using *geomorph* (Adams and Otárola-Castillo 2013). We split each body region into sets of a priori evolutionary modules *(sensu* Klingenberg 2008) roughly aligned with anatomical axes. Head landmarks were assigned to two potential module configurations, one along the anterior/posterior axis (hereafter: A/P), and one along the sagittal/lateral axis (hereafter: S/L; see Figure S2). The A/P grouping separates the anterior (clypeus) area which is related to the feeding apparatus from the posterior of the head which houses the brain and mandible muscles. The D/V axis separates structures more toward the midline of the head (central clypeus, antennae) from the sides (eyes, occipital lobes). Mesosoma landmarks were also assigned to three potential groupings, one along the anterior/posterior axis with bias towards the anterior (hereafter: A/p), one along a similar axis with bias towards the posterior (a/P), and one along the dorsal/ventral axis (D/V; see Figure S2). These anterior/posterior groupings correspond to landmarks associated with different body segments, while the D/V grouping associates landmarks in the region closer the legs or dorsal part of the body, respectively. In this framework, we compared the covariance ratio (CR; Adams 2016) of each hypothesized set of landmarks to those of simulated sets of landmarks (averaged between orientations rotated up to 90° in 0.05° increments), while accounting for phylogenetic relationships. Each simulation test was run for 1000 iterations (see online supplement).

## Results

### Evolutionary Rate

In comparisons of different body regions of the same worker caste, mesosoma shape evolved more rapidly than head shape in both major workers (rate ratio rr = 6.02, p < 0.01) and minor workers (rr = 6.14, p < 0.01; Figure 4). In comparisons of similar traits between worker castes, we observed no significant differences in evolutionary rate for head shape (rr = 1.07, p = 0.59) or mesosoma shape (rr = 1.10, p = 0.59). In contrast to the rate variation among shape traits, evolutionary rates estimated for size traits showed few differences between worker castes or between the head and mesosoma (Figure S4), with the exception of the major worker’s head which evolved relatively slowly.

**Figure 4:**
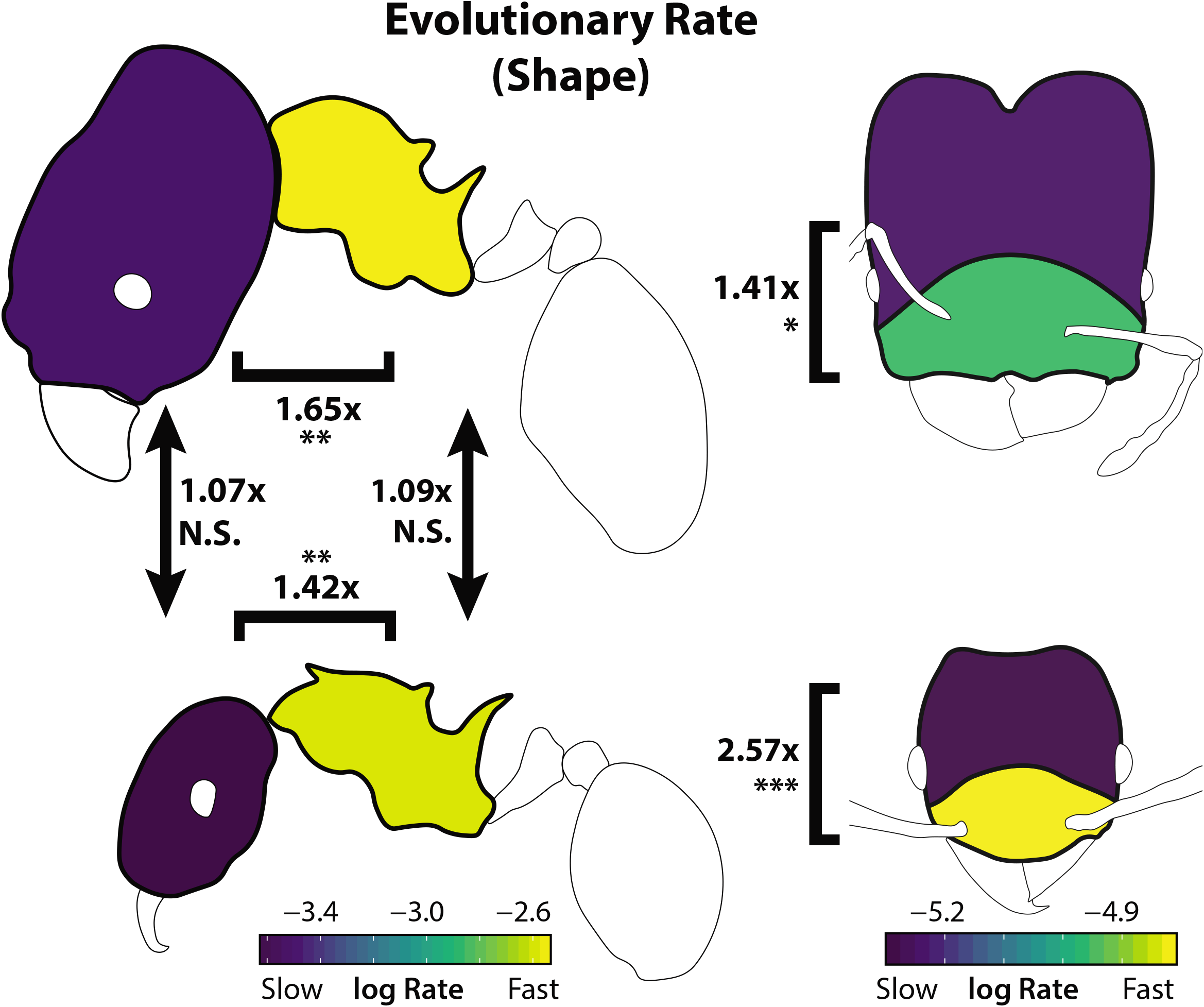
Evolutionary rates are displayed here as a morphogram heat map (Martin & Wainwright 2011). Comparison ratios between traits digitized using different numbers of landmarks (e.g., head and mesosoma) are given as ratios compared to a simulated null ratio. Arrows and brackets indicate statistical tests of rate differences compared to a simulated null, with accompanying numbers describing the estimated rate ratio for the two traits. *p < 0.05, **p<0.01, ***p<0.001

Our tests of modularity within body regions suggested the presence of two evolutionary modules in the *Pheidole* head, in an anterior-posterior arrangement, though the use of semi-landmarks may bias this result (see online supplement). We compared evolutionary rate between the inferred modules of head shape (Figure S2). In these analyses, the anterior landmarks exhibited a higher rate of evolution than the posterior landmarks in both major workers (Figure 4; rr = 1.42, p < 0.05) and minor workers (rr = 1.42, p < 0.001). We also tested for variation in evolutionary rate among lineages. This analysis showed significant support for differences in evolutionary rate of minor workers’ mesosoma shape between biomes, with the most rapid evolution seen in the tropics (see online supplement).

### Morphological Integration

Morphological integration is described here as correlated evolution between morphological shape characters. The strength of this correlation is described using the PLS correlation coefficient (r-PLS), and its significance is assessed by comparison to a simulated null distribution (Adams and Felice 2014; Adams and Collyer 2016). For estimates of body size rather than shape it is measured as the correlation coefficient of independent contrasts (r-PIC).

We found strong indications of morphological integration between both worker castes and body regions in *Pheidole*, however the strength of these correlations varied depending on the comparison (Figure 5a). Head shape was correlated with mesosoma shape in both major workers (r-PLS = 0.53, p < 0.001) and minor workers (r-PLS = 0.51, p < 0.001). In examinations of morphological integration between worker castes, mesosoma shape was strongly correlated between castes (r-PLS = 0.76, p < 0.001), whereas head shape showed a weaker albeit still significant correlation (r-PLS = 0.48, p < 0.001). This difference in worker caste integration effect among body regions was highly significant (two-sample z test; p < 0.001). Similar results were observed for analyses performed with semi-landmarks from the head’s posterior lateral lobes included. Morphological integration varied somewhat between clades (see online supplement), with the Asian-African clade exhibiting a lower degree of integration for all shape traits.

**Figure 5:**
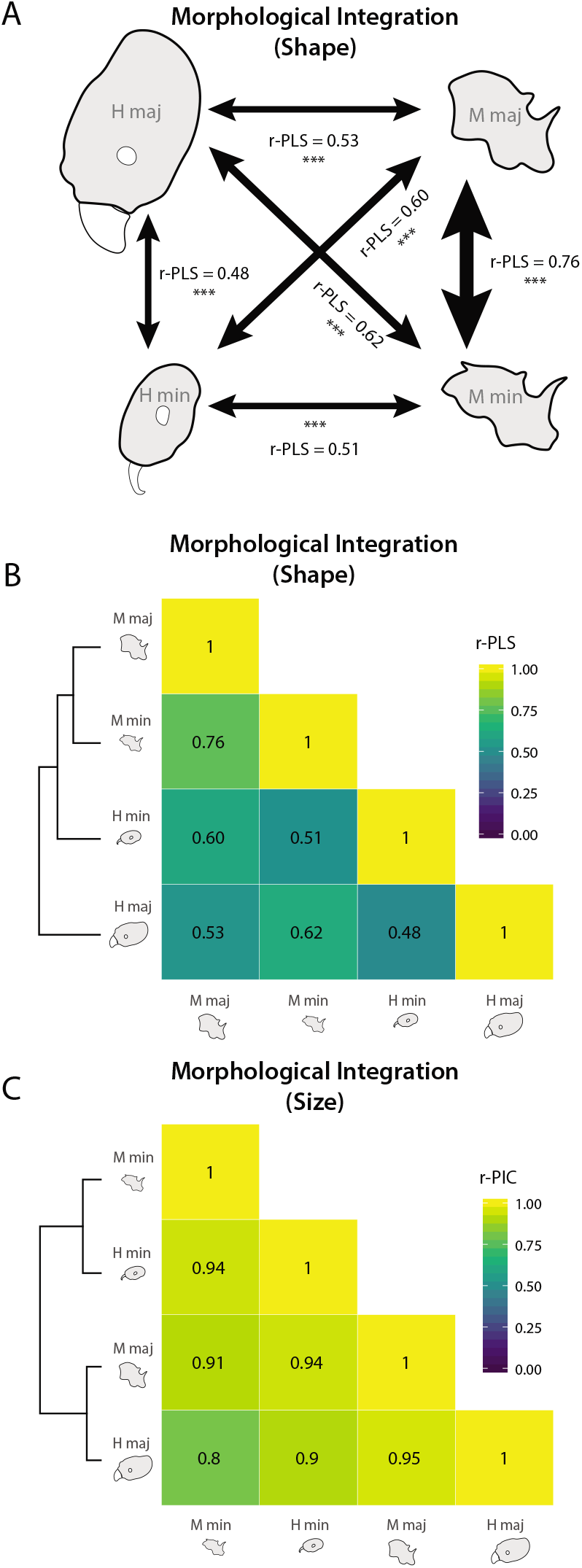
Morphological integration between among body parts within and among worker castes is shown by arrow width in (A). Hierarchical clustering of integration relationships for trait shape is shown in (B) and for trait size in (C), with the strength of relationships indicated by the heatmap and displayed value – r-PLS for trait shape and r-PIC for trait size. *p < 0.05, **p<0.01, ***p<0.001

We performed hierarchical clustering on correlation coefficient matrices for shape integration and size integration (Figure 5b, Figure 5c). Overall, *Pheidole* showed much greater morphological integration in size than in shape. Morphological integration was greater for size traits (r-PIC 0.8 – 0.95) than for any shape traits (maximum r-PLS = 0.76). This integration in size was greater within worker castes than between them (Figure 5c). Morphological integration of shape traits was greatest between the mesosoma of major and minor workers, which evolved as though it were a single module. Head shape was weakly integrated with other traits for minor workers, and least integrated for major workers (Figure 5b).

The scaling relationship between the sizes of different parts is a common theme in evolution and development. As expected, we found a tight relationship between mesosoma size (Weber’s length; Weber 1938) and head length; this was evident in both majors and minors (Figure S3). Relationships between the shape of the head and mesosoma and body size were observable, however they were very weak and poorly predictive (all R-squared values < 0.03; Table S3).

## Discussion

Our results showed varying evolutionary rates and degrees of evolutionary integration within and among worker castes. Overall, evolutionary rate and integration this followed the predictions of different hypotheses (Figure 1). In particular, the mesosoma exhibited integration among homologous traits in different worker castes, while the head exhibited a weaker degree of integration. We found that the mesosoma evolved faster than the head and with a greater degree of morphological integration between castes (Figure 4), but in general evolutionary rate was similar for homologous traits in different worker castes. We found a complex pattern whereby the evolution the head shape of major workers was largely decoupled from that of other traits, but was not necessarily evolving faster.

The evolutionary rate of carapace shape was highly divergent across the different parts of the ant (head vs. mesosoma). This observation was most evident with regards to mesosoma shape, which evolved roughly 1.5x faster than head shape (when corrected for variance differences). Moreover, we found that the anterior portion of the head near the mandibles and mouthparts is evolving more quickly than the posterior half. However, there were no significant differences in evolutionary rate among homologous traits between majors and minors. Thus, homologous traits, and not traits within a caste, tended to evolve at similar rates (Figure 1).

The inferred rate similarity among traits does not alone imply the traits themselves are correlated in their evolution (i.e. they could be evolving at similar rates but on different trajectories), thus we also investigated which sets of traits were correlated during evolution. Here, we found a different pattern, whereby the evolution of mesosoma shape was tightly linked across major and minor workers, but head shape was more decoupled between the two castes. In this way, the head of the major worker was the least integrated with other traits, and the mesosoma of the worker was the most integrated. Previous research in *Pheidole* found that integration among linear measurements was weaker for minor workers than major workers (Pie and Traniello 2007). In contrast, our analyses found weaker integration between head and mesosoma shape for majors than minors. Thus, no one integration hypothesis was supported – either between homologous traits, or between traits within a caste – but rather a mixture of the two.

The fact that mesosoma shape evolved more rapidly than head shape is somewhat surprising, as the head would presumably be the most related to feeding ecology, a key trait that varies across ant species. One potential explanation is that head shape is under stronger stabilizing selection. However, another potential conclusion is that fast mesosoma evolution reflects relative size and arrangement variation in the underlying muscles that control load-carrying and locomotion, which could reflect functional differences in how the ant carries, moves, and performs different tasks. The primary axis of mesosoma variation runs from a stocky shape to a more gracile and elongate one, and most changes are happening repeatedly within limited bounds. There is reason to expect that stocky shapes are common in belowground-foraging species, and that more gracile characteristics are associated with aboveground-foraging and associated defensive traits like spines (Weiser and Kaspari 2006; Sarnat et al. 2017). *Pheidole* are known to vary in the extent to which they live and forage in the leaf litter or on vegetation (Mertl et al. 2010), and there could be tradeoffs inherent the designs adapted for moving and foraging on horizontal vs. vertical surfaces. This would also explain why major and minor mesosomas are tightly integrated in shape, because they face similar biomechanical challenges due to living and moving in similar environments. Thus, these phenotypes may represent ecomorphs that are repeatedly evolved in each newly colonized region, as in *Anolis* lizards (Mahler et al. 2013). However, given the paucity of behavioral observations for most ant species around the world, further study is required to understand this trait’s functional and biomechanical significance. Furthermore, the linking of external geometry with variation in underlying function and performance remains an important avenue for future work on comparative anatomy and biomechanics in ants.

We find support for the hypothesis that the shape of minor and major worker castes can evolve independently (Holley et al. 2016), promoting the evolution of ecological specialization. We emphasize that this is not simply a statement that head shapes are different between majors and minors, which is obvious, but that they can evolve on diverging trajectories (i.e. the major is not just a consistent transformation of the minor). This allows for increased evolutionary “degrees of freedom” in the functional specialization among castes. However, this finding was specific to the head region, as mesosoma shape was tightly integrated across castes. The fact that rates of shape evolution were 1.5 times greater for the highly integrated mesosoma than for the head (Figure 4) suggests that integration in this case does not constrain, but may rather accelerate rates of evolutionary divergence in shape among species (Cheverud 1995; but see Márquez and Knowles 2007).

Allometry is a common theme and pattern in development and evolution, and strong relationships between the sizes of different body parts are expected during evolution. Matching this expectation, we found that head and mesosoma sizes were tightly linked both within and among castes (Figure 5). In contrast to the pattern for cranial evolution in birds (Klingenberg and Marugán-Lobón 2013), relationships between shape traits and body size were significant, but poorly predictive (Figure S3; Table S1). While we were not able to account for allometric relationships within species due to our study design, we did find that cross-species relationships between body size and shape traits were not strong enough to potentially drive other patterns reported in this study. Our estimates of evolutionary rate for size traits showed that the size of each trait evolved faster than its shape (Figure S4), confirming a similar observation by Pie and Tscha (2013). Interestingly, major worker heads evolved at the slowest rate for size and among the slowest for shape despite being the least integrated with other body parts (which should thus release it from constraint by pleiotropic effects; but see Cheverud 1995). This suggests that this trait is more evolutionarily conserved; future studies investigating the evolutionary consistency of major worker tasks (as in Mertl et al. 2010) and their biomechanical needs would be valuable in explaining this pattern.

In principle, correlations in size and shape among traits/castes could be caused by either selection or developmental constraint. This kind of comparative analysis does not by itself allow for inference of the underlying selective or developmental mechanisms responsible for the patterns of integration that we identify. However, there is a strong body of work on the developmental basis of caste differentiation in *Pheidole*, and especially the role of JH as a developmental switch cues, that can inform the likelihood of some potential explanations. Notably, classic (Wheeler and Nijout 1983, 1984; Wheeler 1991) and more recent (Rajakumar et al. 2012; Lillico-Ouachour and Abouheif 2016) work shows that experimental manipulation of pheromone exposure can alter the relative sizes of *Pheidole* majors and minors, and manipulation of rudimentary wing discs can alter the relative sizes of the head and body (Rajakumar et al. 2018). Moreover, in other insects, it has been shown that relative sizes of different body parts can be experimentally selected for (Frankino et al. 2005; Stillwell et al. 2016). If researchers can manipulate relative size with apparent ease using chemical cues or artificial selection, this implies that evolution may not be constrained from doing the same. We expect that general diversification of body size is likely to due to selection on loci that control body size overall, rather than independent selection on the size of each part. However, the fact that relative sizes of different parts have been maintained in evolutionary time implies selective advantages of the relative sizes of body parts within and among castes (Gould 1966).

To our knowledge, less is known about the developmental basis of the shape characters we are capturing in our landmark system, so developmental constraints or biases may explain some of the evolutionary correlation in shape we observe. However, the evolutionary modules in the head inferred by our analysis (Figure S2) do not correspond to the head developmental modules inferred by Yang and Abouheif (2011) in their examination of *Pheidole* gynandromorphs. If both studies are correct, this would imply that developmental modularity does not underlie the macroevolutionary modularity we infer, leaving selection and non-genetic influences, as well as methodological issues with comparing fixed landmarks and semi-landmarks, as the most likely explanations for why different regions of the head appear to evolve separately or independently. An interesting future direction would be to attempt to experimentally investigate the developmental bases of the axes of shape variation we identify in our study.

One noticeable feature of the genus *Pheidole* s global diversification has been the re-evolution of similar environmental and behavioral niches in different geographic regions, each radiation following from a single colonization event (Moreau 2008; Economo et al. 2015a). While morphological evolution in this clade has been largely conserved throughout its history (Pie and Traniello 2007), similar body size phenotypes have consistently re-evolved following each clade’s colonization of a new biogeographic realm (Economo et al. 2015a). In this study we observed that New World and Old World radiations of *Pheidole* occupied mostly overlapping portions of morphospace (Figure 3), whereas the Australasian clade occupied a smaller, but still overlapping portion of this same trait space. We found this pattern for size and shape of both head and mesosoma. It remains unclear why some portions of morphospace, and large body size in particular, have not evolved in Australasian taxa. One potential explanation is that niche filling in this most recent radiation is ongoing – indeed the Australian clade is the youngest of the continental radiations and is still in a more elevated phase of its diversification (Economo et al. 2019).

## Conclusion

The morphological and functional differentiation of castes is thought to be a key evolutionary innovation underlying the success of ants and other social insects. Patterns of macroevolutionary integration and modularity within and among castes may provide clues to the selective forces shaping diversification in ants, and the developmental biases and constraints involved in trait divergence (West-Eberhard 1979). We find that size evolution is tightly integrated and evolving with homogeneous rates both among parts in a single caste, and across the worker castes. In contrast, our results using geometric morphometric estimates of body shape indicate that while mesosoma shape shows homology integration, head shape has become largely disintegrated between major and minor workers (Figure 3c). Head morphology and its associated musculature is associated with ecological specialization in many taxa, often but not exclusively due to feeding functionality, thus the differences in head shape between major and minor workers probably represent divergence in their tasks in the colony (Smith 1987; Futuyma and Moreno 1988; Mertl and Traniello 2009). In this case, evolution of developmental pathways facilitating independent evolution of major and minor worker phenotypes could represent key innovations enabling lineages with this trait to occupy multiple specialized strategies at once, or to discover new team strategies emergent from their polymorphism (Wheeler & Nijhout 1981, 1984; Wheeler 1990; Anderson and McShea 2001). Interestingly, the independent evolution of the head does not lead to faster rates of evolution, and in fact mesosoma shape evolves 1.5x faster than head shape in *Pheidole*. We hypothesize that this rapid evolution of the mesosoma reflects a pattern of frequent adaptation to different biomechanical needs in different microhabitats, but future work is needed to test this hypothesis.

While body-size polymorphism is a common trait in ants, “complete” polymorphism (i.e., in shape) is rarer but noticeably present in some of the most diverse ant clades (Wills et al. 2017), an observation that hints at a role for polymorphism in adaptability (Wilson 2003). We propose that, beyond the benefits of body-size polymorphism, the reduction of morphological integration between distinct behavioral strategies, inclusive of sexes, castes, and alternative reproductive tactics (West-Eberhard 1979), could be a recurring key innovation that enables the evolution of adaptive polymorphism and promotes rapid diversification. Further comparative studies on the evolution integration and modularity across radiations of ants with worker polymorphisms, and any concurrent changes in diversification rates and patterns, would be useful for testing this hypothesis.

## Acknowledgements

We thank R. Keller, Y. Hashimoto, C. Peeters, S. Price, A. Saurez, and members of the Economo Unit for providing stimulating discussion, and A. Lazarus and M. Ogasawara for photographing specimens. C. Klingenberg and several anonymous reviewers provided constructive comments that contributed to this work. EPE, LLK, and JPH were supported by NSF (DEB-1145989). NRF, BL, GF, EMS, and EPE were supported by subsidy funding to OIST, and EPE and NRF were supported by Japan Society for the Promotion of Science KAKENHI grants (17K15180 and 17K15178, respectively).

