## Supplementary material for "Macroevolutionary integration of phenotypes within and across ant worker castes": Online Supplement

### Rate Heterogeneity across Lineages

We tested whether rates of trait evolution differed depending on assignment of lineages to three a priori groupings: clade, biome, and diet (Adams 2014). To define these groups, we used geographic coordinates from our own compilation including collection records of the specimens contributors, supplemented with data from AntWeb.org, to associate geographic and environmental data with each specimen. We assigned species into “temperate” and “tropical” categories using latitude data, applying a majority rule to species with specimens associated with multiple categories. We used a similar approach in assigning species to one of 4 biome categories (A - D) following the Köppen classification scheme (Köppen 1884). In this framework, A refers to moist tropical climates, B refers to dry climates, C refers to humid coastal/subtropical climates, and D refers to interior continental climates. Biome classification for each set of coordinates was extracted from publicly available raster data based on the WorldClim climate dataset (Hijmans et al. 2005). The data on granivory is mostly derived from Moreau (2008), with additions of field data for a few Old World species (see online supplement).

Species in the three major *Pheidole* radiations exhibited no significant differences in rates of evolution for head shape (Table S1; majors: rr_max_ = 1.10, p = 0.97; minors: rr_max_ = 1.32, p = 0.63). However, mesosoma shape evolved more rapidly in the Australia/PNG clade for majors (rr_max_ = 1.88, p < 0.39), and significantly more rapidly in both the Australia/PNG and New World clades in minors (rr_max_ = 2.67, p < 0.01).

Lineages confirmed to exhibit granivorous behavior (Moreau 2008) showed no significant differences in rates of head or mesosoma evolution compared to other lineages. While rates of mesosoma evolution were greater in granivorous species, this difference was not significant (Table S1; majors: rr = 1.76, p = 0.63; minors: rr = 2.47, p = 0.36).

Comparing rates of trait evolution between biomes showed no significant differences for head shape, however there were major differences in the rate of mesosoma shape evolution in minor workers (rr_max_ = 9.13, p < 0.05; majors: rr_max_ = 2.78, p = 0.80): in this model, species in equatorial and warm continental biomes evolved more rapidly. Lastly, no significant rate differences were observed between temperate and tropical lineages for either head shape (majors: rr = 1.08, p = 0.71; minors: rr = 1.18, p = 0.40) or mesosoma shape (majors: rr = 1.13, p = 0.61; minors: rr = 1.35, p = 0.19).

We also repeated our rate estimation analyses independently for each clade. Generally, we estimated slower evolutionary rates of size evolution for the New World clade than for the Old World or Australasian clades (Figure S1). However, this pattern was not observed for shape traits, for which the Australasian clade showed the fastest evolutionary rates for mesosoma shape, and the Old World clade showed the slowest for this trait.

A similar comparison based on individual *Pheidole* clades was also conducted for morphological integration among traits. In this analysis, we found that size was most integrated the Old World clade, followed by the Australasian clade and the New World clade in every case (Figure S2). In contrast, Old World *Pheidole* showed weaker integration for shape traits. Despite these differences, the greater integration of mesosoma shape between worker castes in comparison to head shape was consistent across clades.

### Modularity Within Body Regions

Modularity is measured in *geomorph* as the covariance ratio (CR), which describes the covariation between modules relative to the covariation within modules (Adams 2016). When CR ≥ 1, modules show no observable signal of modularity; values significantly less than 1 (compared to a simulated null distribution) indicate independence between modules. We estimated modularity for two a priori configurations for head landmarks, and three a priori configurations for mesosoma landmarks (Figure S1).

We tested for modularity in head shape using two datasets: one including only fixed landmarks, and one also including semi-landmarks describing the curvature of the posterior lateral lobes. Using only fixed landmarks, head shape showed no significant modularity along the A/P axis in major workers (CR = 1.31, p = 1.00; see also Figure S4) or minor workers (CR = 1.28, p = 1.00). We observed similarly non-significant scores when dividing landmarks into a S/L axis; this was consistent across both major workers (CR = 1.17, p = 0.91) and minor workers (CR = 1.18, p = 0.87). However, when we included (semi-landmark) data on the posterior lateral lobes, we observed that head shape evolved as two independent modules along the anterior/posterior axis in major workers (CR = 0.79, p = 0.003) and in minor workers (CR = 0.76, p = 0.002). No such effect was observed along the sagittal/lateral axis for either major workers (CR = 0.98, p = 0.26) or minor workers (CR = 1.05, p = 0.69). For mesosoma shape, we observed no significant or even weak evidence of modularity (always CR > 1; see Figure S4). However, few potential module configurations exist for a set of only 6 landmarks, which likely limited our ability to detect modularity with this dataset.

We observed significant modularity in head shape for both minor workers and majors when dividing the head into anterior and posterior modules (Figure S4). Among these two modules, we found that the anterior evolved between 1.5 and 2.5 times faster than the posterior module when correcting for the difference in number of landmarks. This result suggests that the anterior portion of the head has evolved more rapidly, potentially as species’ functional traits have adapted to different lifestyles in each radiation (Wilson 2003; Economo et al. 2015). While many of the more dramatic differences among *Pheidole* species are concentrated in the posterior end of the head – for example with rounded vs. extended lobes (Fischer et al. 2012) – these may be more weakly affected by selection during diversification. Alternatively, while the posterior lobes have muscle attachment points for the mandibles, and indeed are often extended in lineages with exceptional mandible strength (Paul and Gronenberg 1999), they may be a part of a more conserved evolutionary module whose change might require rearrangements of complex internal structures, as in the avian cranium (Klingenberg and Marugán-Lobón 2013). Some caution is advised in interpreting these results however, as the use of semi-landmarks exclusively in the posterior margin of the head may under-estimate variation among species in this region, and thus inflate the differences observed between the posterior and anterior regions of the head.

While we found no indication of modularity in mesosoma shape, we did observe that the anterior region of the mesosoma (a/P configuration in Figure S1) evolved more rapidly in both subcastes. Variation in this region appears to be associated with size changes in the pronotum, and so may indicate functional changes in the musculature responsible for lifting the head (Keller et al. 2014; Sarnat et al. 2016). In the standard photographic views we used, the ant’s head obscured the outline of the anterior pronotum and thus prevented our measuring it from photographs. This is the segment that includes most muscles involved in supporting the head (Keller et al. 2014), and thus we may have missed some variation in mesosoma shape that should be mechanistically linked to head shape. Future studies of morphology and modularity in particular, will undoubtedly benefit from the use of 3D scanning and morphometric approaches, which allow the measurement of traits typically obscured in photographs (Hita Garcia et al. 2017).

### Supplemental Figures

#### Figure S1: Rate differences between clades. Each analysis follows methods described for analysis of the complete dataset, but was performed on a subset of the data.

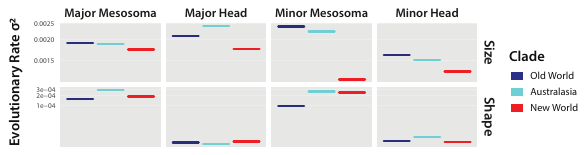

#### Figure S2: Comparison of morphological integration estimates among clades. Each analysis follows methods described for analysis of the complete dataset, but was performed on a subset of the data. The lower triangle shows results from the complete dataset, and the upper triangle includes bars indicating the correlation coefficient (r-PLS or r-PIC) estimated for each clade.

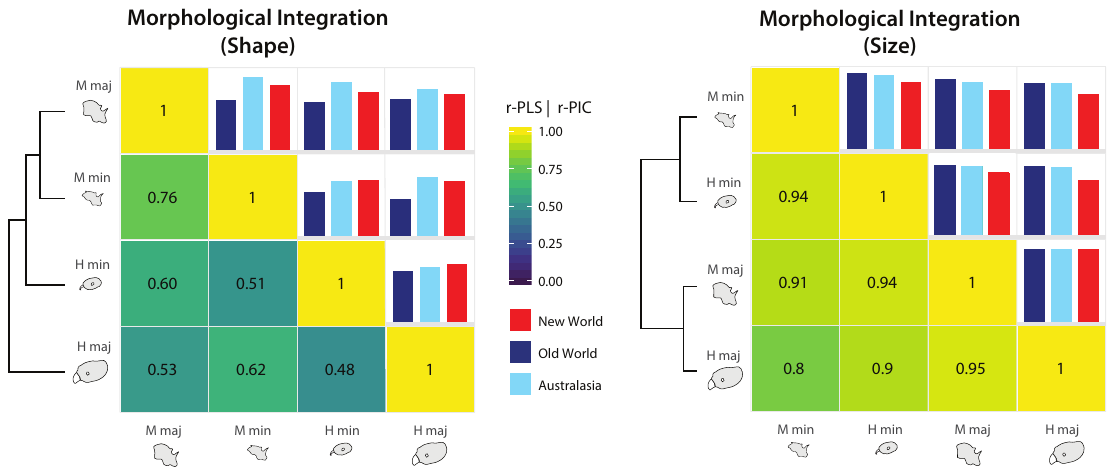

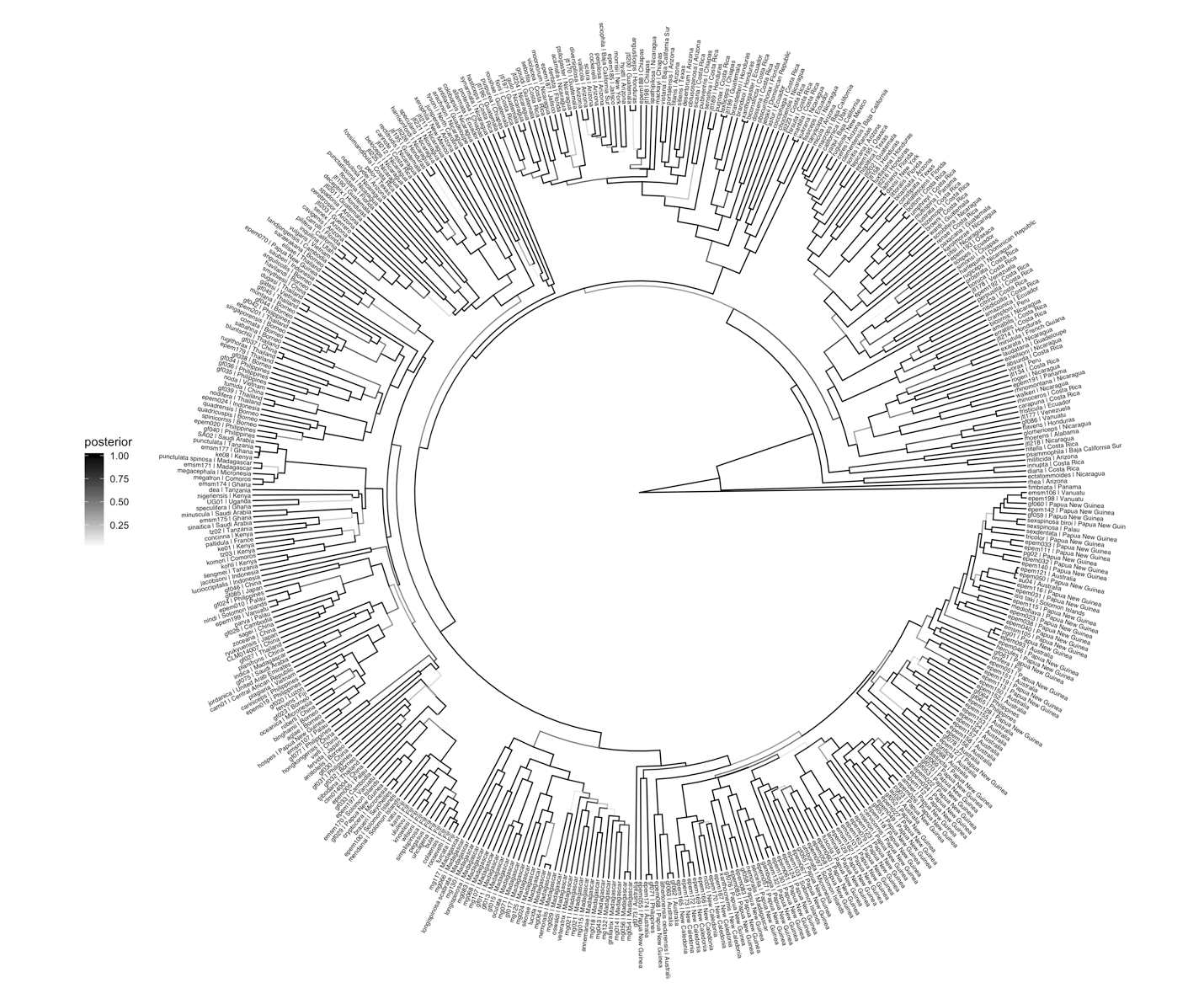

#### Figure S3: Phylogeny used in this study (Economo et al. 2019). Posterior clade probabilities are displayed as branch colors (see legend at left).

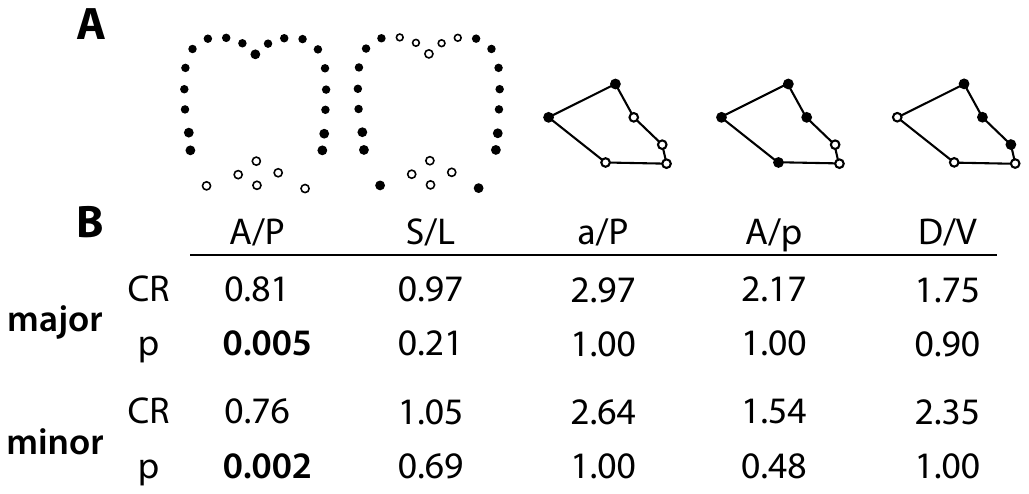

#### Figure S4: Hypothesized evolutionary module configurations (A) are shown with landmarks assigned to each module assigned different colors. The table in (B) shows results of phylogenetically-corrected modularity tests conducted in geomorph. Covariance Ratios (CR) are given for each hypothesized configuration, as well as p-values derived from comparison against a simulated null.

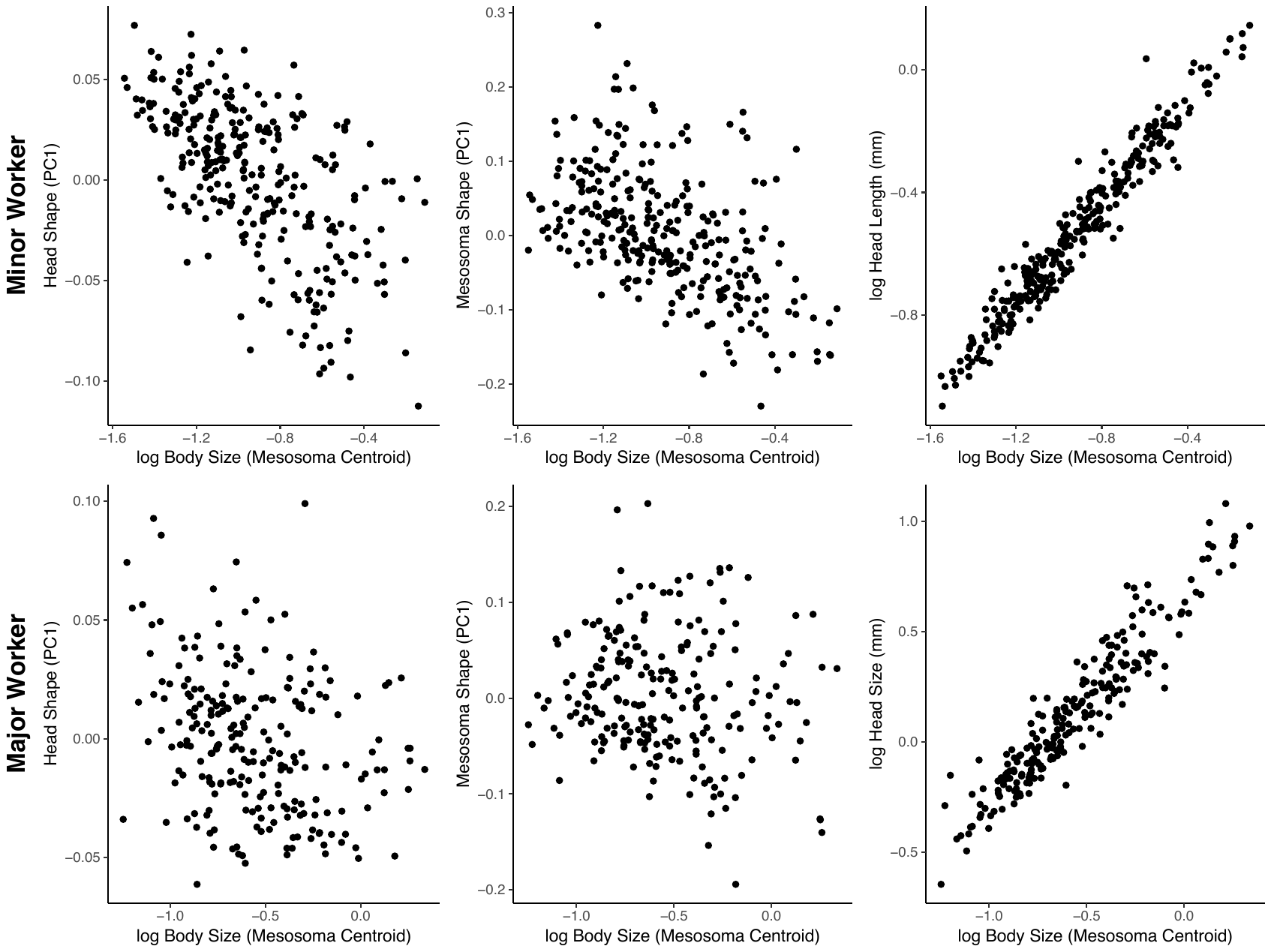

#### Figure S5: Scatter plot showing species averages, comparing the first principle component of head (left column), mesosoma shape (center column) and log-transformed body size. Minor workers are shown in the top row, and major workers are shown in the bottom row. The right column shows the standard allometric relationship between head length and body size.

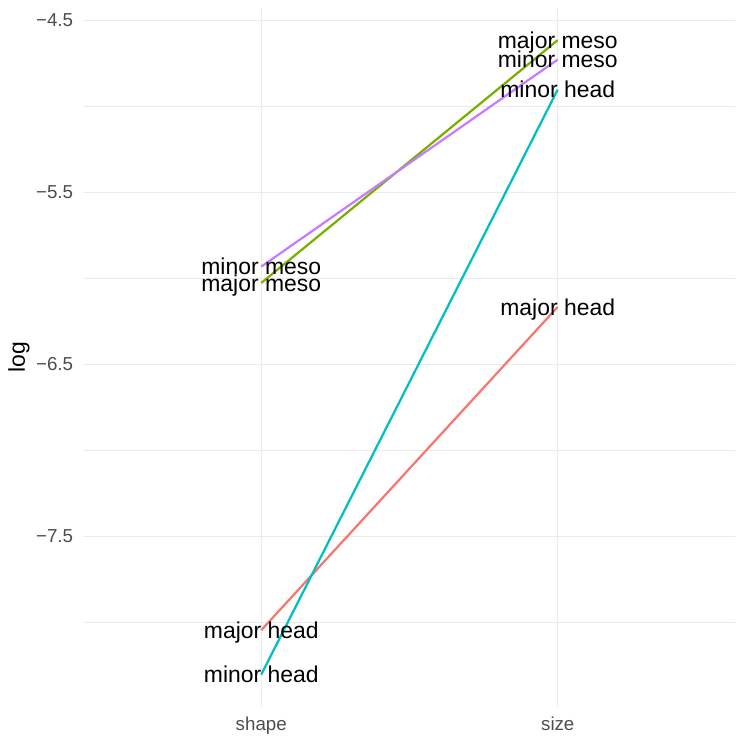

#### Figure S6: Comparison of evolutionary rate estimates for size and shape of *Pheidole* body parts and worker castes.

#### Table S1: Landmark definitions for *Pheidole* head.

| **Number** | **Type** | **Definition** |
| --- | --- | --- |
| **1** | Fixed | Left eye – Posterior apex |
| **2** | Fixed | Left eye – Anterior apex |
| **3** | Fixed | Right eye – Posterior apex |
| **4** | Fixed | Left eye – Anterior apex |
| **5** | Fixed | Clypeal median – Posterior margin apex |
| **6** | Fixed | Clypeal median – Anterior margin apex |
| **7** | Fixed | Left mandible intersection with lateral clypeal margin |
| **8** | Fixed | Right mandible intersection with lateral clypeal margin |
| **9** | Fixed | Left antenna torulus – Anterior margin apex |
| **10** | Fixed | Right antenna torulus – Anterior margin apex |
| **11** | Fixed | Occipital center – Posterior Apex |
| **12 – 19** | Semi | Seven equidistant points along left occipital margin between landmarks 3 and 1, with right side removed |
| **20 – 25** | Semi | Reflection of points 12 – 19. |

#### Table S2: Landmark definitions for *Pheidole* mesosoma.

| **Number** | **Type** | **Definition** |
| --- | --- | --- |
| **1** | Fixed | Propodeal insertion of petiole – Superior |
| **2** | Fixed | Propodeal insertion of petiole – Inferior |
| **3** | Fixed | Propodeal spiracle |
| **4** | Fixed | Intersection of mesopleural, propodeal margins – Inferior apex |
| **5** | Fixed | Intersection of pronotal, mesopleural margins – Inferior apex |
| **6** | Fixed | Intersection of mesopleural, propodeal margins – Superior apex |

#### Table S3: Results of phylogenetic regressions between body size (measured as the centroid size of the mesosoma) and shape traits.

| **Trait** | **z** | **R^2^** | **p** |
| --- | --- | --- | --- |
| Minor head shape | 1.994 | 0.01 | 0.018 |
| Major head shape | 3.005 | 0.03 | 0.001 |
| Minor mesosoma shape | 1.358 | 0.01 | 0.090 |
| Major mesosoma shape | 2.580 | 0.03 | 0.007 |

| Table S4: Rates of trait evolution (log_10_) across clades defined by continental expansions and climatic regimes | | | | |
| --- | --- | --- | --- | --- |
|  | **Major** | | **Minor** | |
| **Clade** | **Head** | **Mesosoma** | **Head** | **Mesosoma** |
| New World | -5.06 | -3.72 | -5.10 | -3.60 |
| Old World | -5.10 | -3.80 | -5.06 | -4.00 |
| Australasia | -5.14 | -3.53 | -4.93 | -3.57 |
| **p** | 0.97 | 0.41 | 0.62 | **0.004** |
|  | **Major** | | **Minor** | |
| **Biome** | **Head** | **Mesosoma** | **Head** | **Mesosoma** |
| Tropical (A) | -5.08 | -3.65 | -4.99 | -3.69 |
| Dry (B) | -5.23 | -3.98 | -5.00 | -4.10 |
| Temperate (C) | -5.13 | -3.66 | -5.13 | -3.50 |
| Continental (D) | -5.32 | -4.09 | -5.23 | -4.46 |
| **p** | 1.00 | 0.78 | 1.00 | **0.03** |
|  | **Major** | | **Minor** | |
| **Latitude** | **Head** | **Mesosoma** | **Head** | **Mesosoma** |
| Temperate | -5.12 | -3.73 | -5.07 | -3.59 |
| Tropical | -5.09 | -3.68 | -5.00 | -3.72 |
| **p** | 0.71 | 0.56 | 0.40 | 0.20 |
